# Reproduction and longevity A Mendelian randomization study of gonadotropin-releasing hormone and ischemic heart disease

**DOI:** 10.1101/472548

**Authors:** C M Schooling, Jack C M Ng

## Abstract

**Background:** According to well-established evolutionary biology theory reproduction trades-off against longevity, implying that upregulating the reproductive axis might drive major diseases. We assessed whether the central driver of reproduction gonadotropin-releasing hormone 1 (GnRH1) had a causal effect on the leading cause of global morbidity and mortality, i.e. ischemic heart disease (IHD). As a contrast we similarly examined the role of GnRH2 because it is more a driver of female sexual behavior.

**Methods:** We applied strong (p-value <5×10^−6^) and independent genetic predictors of GnRH1 and GnRH2 to an extensively genotyped IHD case (n=76,014) - control (n=264,785) study using multiplicative random effects inverse variance weighted (IVW), weighted median, MR-Egger and MR-PRESSO estimates.

**Results:** GnRH1, predicted by 11 genetic variants, was positively associated with IHD (IVW odds ratio (OR) 1.04 per effect size, 95% confidence interval (CI) 1.01 to 1.08), but GnRH2, predicted by 15 genetic variants, was not (IVW OR 0.98, 95% CI 0.95 to 1.02).

**Conclusions:** GnRH1 is a potential IHD genetic target. Apart from demonstrating a central tenet of evolutionary biology in humans, our study suggests that existing treatments and environmental factors targeting GnRH1, its drivers or consequences could be re-purposed to prevent and treat IHD. Given, the importance of reproduction to the human species, many such exposures likely exist.

## Background

Despite substantial progress in prevention and control, cardiovascular disease (CVD) remains incompletely understood.^1^ CVD reduction is vital to achieving the Sustainable Development Goals.^2^ To address this gap, CVD is being re-conceptualized within the well-established evolutionary biology theory that growth and reproduction trade-off against longevity.^3 4^ Dietary restriction mimetics are being investigated in CVD.^5^ Genetic selection in favour of both fertility and ischemic heart disease (IHD) has been demonstrated.^6^ Reproduction is driven centrally by gonadotropin-releasing hormone 1 (GnRH1)^7^ likely driven by environmental factors to ensure optimal timing of reproduction. We used Mendelian randomization to estimate the causal effect of GnRH1 on IHD. Mendelian randomization uses genetic proxies of exposure. Genetic make-up is determined randomly at conception giving randomization akin to that in a randomized controlled trial,^8^ and thereby unconfounded estimates. We also considered GnRH2, because it is more a driver of female sexual behavior than reproduction,^7 9 10^ so GnRH1 would be expected to have more of an effect on IHD.

## Methods

### Gonadotropin-releasing hormone 1 and 2

Genetic associations with GnRH1 and GnRH2 protein were obtained from a genome-wide association study (GWAS) (n=3,301; mean age, 43.7 years; 1,615 women [49%]) of people of European descent.^11^ Genetic associations with rank inverse normalized residuals were obtained from linear regression of natural log-transformed protein abundances adjusted for age and sex, duration between blood draw and processing and the first three principal components of ancestry from multidimensional scaling to account for population stratification.^11^

We used all single nucleotide polymorphisms (SNPs) which independently (r^2^<0.05) and strongly (p-value <5 × 10^−6^) predicted GnRH1 or GnRH2. We used the “clump_data” function of MRBase^12^ to select between SNPs in linkage disequilibrium. We assessed the strength of genetic instruments from the F-statistic using an approximation.^13^ We discarded SNPs associated with potential confounders, including current smoking, age at completion of full-time education, Townsend index, alcohol intake frequency and number of days per week walked for 10+ minutes, at Bonferroni corrected significance in the UK Biobank GWAS summary statistics (https://github.com/Nealelab/UKBiobankGWAS) of ≤361,194 people of White British ancestry. To assess whether the SNPs might affect IHD other than via GnRH we searched two curated genotype to phenotype databases, Ensembl release 94 (http://useast.ensembl.org/index.html) and PhenoScanner,^14^ for paths by which the SNPs predicting GnRH1 or GnRH2 might affect IHD other than via GnRH. We searched PhenoScanner for diseases and traits associated with the SNPs (or their correlates r^2^>.0.8) at p-value<lxl0^−5^.

### Ischemic heart disease

Genetic associations with IHD were obtained from the largest publicly available extensively genotyped IHD case (n=up to 76,014)-control (n=up to 264,785) study based on a meta-analysis of the CARDIoGRAPMplusC4D 1000 Genomes case (n=60,801)-control (n=123,504) study/ MIGen/CARDIoGRAM Exome chip study, the UK Biobank SOFT CAD study (cases=10,801, controls=137,371), and two case (n=4120)-control (n=3910) studies from Germany and Greece.^15^ CARDIoGRAMplusC4D 1000 Genomes participants are largely of European descent (77%) with detailed phenotyping based on medical records, clinical diagnosis, procedures that indicate IHD, such as revascularization, and/or angiographic evidence of stenosis, and sometimes case status ascertained from medications or symptoms that indicate angina or from self-report.^16^ In the UK Biobank SOFT CAD GWAS the case phenotype included fatal or nonfatal myocardial infarction, percutaneous transluminal coronary angioplasty or coronary artery bypass grafting, chronic IHD and angina. Controls were defined as individuals who were not a case after exclusions.^15^

### Statistical analysis

SNP-specific Wald estimates were combined using inverse variance weighting (IVW) with multiplicative random effects. Wald estimates were calculated as the SNP on IHD estimate divided by the SNP on GnRH1 or GnRH2 estimate. As sensitivity analysis we used a weighted median, which may provide correct estimates if the SNPs are invalid instruments for <50% of the weight, MR-Egger, which is robust to pleiotropy assuming the instrument strength is independent of the direct effect,^17^ and Mendelian randomization pleiotropy residual sum and outlier (MR-PRESSO), which detects potentially pleiotropic SNPs from the residual sum of squares and removes statistically significant outliers.^18^ Finally, we repeated the analysis using SNPs predicting GnRH1 or GnRH2 at p-value<5 × 10^−8^. We replaced palindromic SNPs by proxies. We used the R packages “MendelianRandomization” and “MRPRESSO” to obtain estimates. We used R (version 3.5.1, The R Foundation for Statistical Computing, Vienna, Austria) for our analysis.

### Ethics

We used publicly available summary data with no direct involvement of participants in the study. No ethical approval is required.

## Results

Of the 225 SNPs predicting GnRH1 at 5 × 10^−6^ one of the 12 independent SNPs (rs145898851) was not available for IHD and had no proxy with r^2^>=0.5, giving 11 SNPs, of which two were replaced by proxies (rs28529564byrsll4767131 (r^2^=0.59) andrsll039086byrs57323109 (r^2^=0.95)). Of the 123 SNPs predicting GnRH2 one invalid SNP (chromosome position, 20:2975929) was discarded. Of the 15 independent SNPs, three were replaced by proxies (rs34246308 by rs1941401 (r^2^=0.98), rs79590491 by rs79238287 (r^2^=0.84), and rs6999411 by rs3019086 (r2=0.50)). The F-statistic for GnRH1 was 33 and for GnRH2 was 24. None of the SNPs were associated with potential confounders in the UK Biobank.

GnRH1 was positively associated with IHD using IVW with little evidence of pleiotropy (Figure 1a). The weighted median, MR-Egger and MR-PRESSO gave similar estimates (Table 1), again with little evidence of pleiotropy. Some genetic predictors of GnRH1 were associated with potential consequences of GnRH1 and causes of IHD (rs141012986 with height and atrial fibrillation, rs2731674 with height, coagulation factors, metabolites, interleukins, lung function, body composition, and rs4241818 with thrombosis and metabolites). GnRH2 was not associated with IHD using IVW with little evidence of pleiotropy (Figure 1b). The weighted median, MR-Egger and MR-PRESSO gave similar estimates with little evidence of pleiotropy (Table 1). Rs34439826 predicting GnRH2 was associated with height and forced vital capacity, while rs79238287 was associated with estrogen treatment.

**Figure 1.**
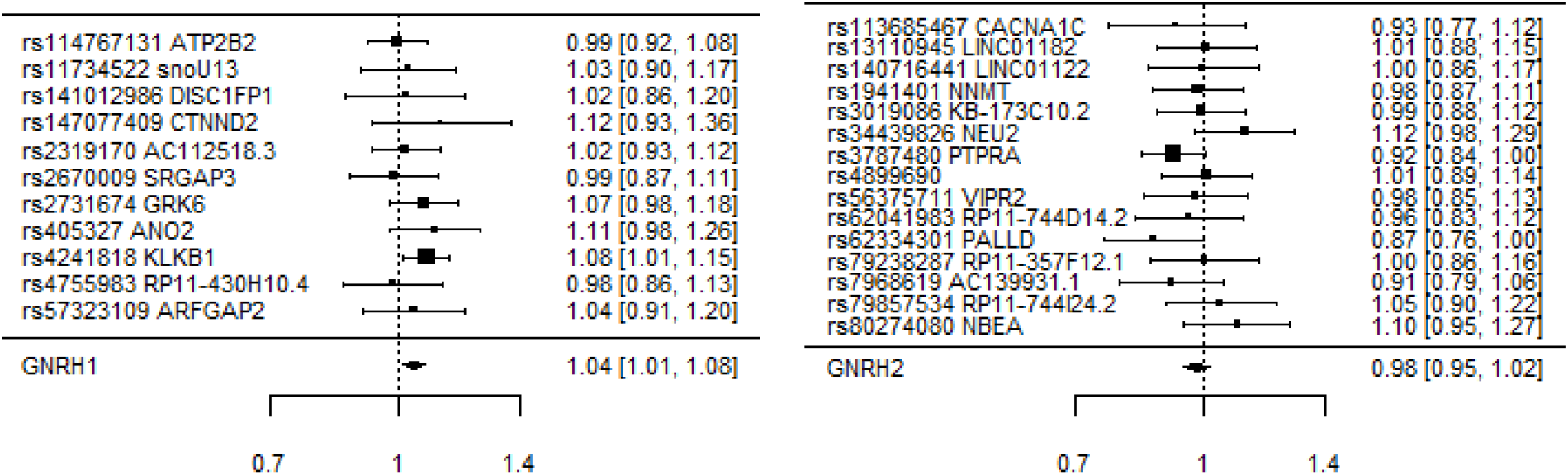
Overall IVW (random effects) and SNP-specific estimates for the association of (a) GnRH1 and (b) GnRH2 with IHD.

**Table 1.** Mendelian randomization estimates for associations of GnRH1 (based on 11 independent SNPs with p-value <5xl0^−6^) and GnRH2 (based on 15 independent SNPs with p-value <5xl 0^−6^) with IHD, using a study largely based on CARDIoGRAPMplusC4D 1000 Genomes and the UK Biobank SOFT CAD.

| Exposure | Mendelian randomization method | Odds ratio | 95% confidence interval | p-value | Cochran's Q statistic (p-value) | MR-Egger |  |
| --- | --- | --- | --- | --- | --- | --- | --- |
|  |  |  |  |  |  | Intercept p-value | I<sup>2</sup> |
| GnRH1 effect size | Inverse variance weighted | 1.04 | 1.01, 1.08 | 0.01 | 6.06 (0.81) | - | - |
|  | Weighted median | 1.04 | 1.00, 1.09 | 0.08 | - | - | - |
|  | MR-Egger | 1.02 | 0.93, 1.13 | 0.61 | 5.92 (0.75) | 0.70 | 75.5% |
|  | MR-PRESSO (corrected) | 1.04 | 1.03, 1.05 | 0.01 |  | - | - |
| GnRH2 effect size | Inverse variance weighted | 0.98 | 0.95, 1.02 | 0.29 | 13.3 (0.50) | - | - |
|  | Weighted median | 0.99 | 0.94, 1.04 | 0.64 | - | - | - |
|  | MR-Egger | 0.98 | 0.88, 1.10 | 0.75 | 13.3 (0.43) | 0.99 | 0.0% |
|  | MR-PRESSO (corrected) | 0.98 | 0.95, 1.02 | 0.30 |  | - | - |

At genome-wide significance, 2 SNPs (rs4241818 and rs2731674) independently predicted GnRH1 and 1 SNP (rs3787480) predicted GnRH2. Based on these two SNPs GnRH1 was positively associated with IHD (IVW with fixed effects odds ratio, 1.08; 95% confidence interval, 1.02 to 1.14). Based on rs3787480 GnRH2 was not associated with IHD (Figure 1b).

## Discussion

Consistent with theoretical predictions from evolutionary biology and empirical evidence,^6^ this study showed GnRH1, but not GnRH2, positively associated with IHD.

This first study assessing the role of GnRH in IHD makes cost-effective use of publicly available GWAS, nevertheless several considerations bear mention. First, although the F-statistics were >10, weak instrument bias is possible, which likely biases towards the null. Second, confounding by population stratification is possible; however both samples mainly concern people of European descent with genomic control,^11 15^ and the SNPs were not associated with potential confounders. Third, some SNPs predicting GnRH1 were associated with factors relevant to cardiovascular disease, however GnRH1 is the master of reproduction^7 9 10^ suggesting vertical rather than horizontal pleiotropy. Fourth, Mendelian Randomization gives unconfounded estimates, but is open to the selection bias that may occur in observational studies of older adults,^19^ because of missing prior deaths from the same underlying cohort. However, IHD occurs at relatively young ages and generally precedes other types of cardiovascular disease, making it less open to such bias. Fifth, GnRH1 and GnRH2 were not measured in the sample with the outcome, however genetic predictors of proteins should not vary by sample. Sixth, effects of genetic variants on IHD could vary by age or sex, but sufficiently large unbiased sex-specific samples are not available to test this possibility. Seventh, we assumed linear associations. Eighth, canalization could generate a bias, however the reproductive axis is most active before birth, during the mini-puberty of infancy and from puberty. Ninth, reproduction trades off against other drivers of longevity, such as immunity,^20^ and so might be expected to promote infections and protect against auto-immune conditions. However, these predications are difficult to test, because currently sufficiently large densely genotyped GWAS of relevant events are not available.

Physiologically GnRH1 drives reproduction,^7 9 10^ including sex hormones. In randomized controlled trials (conjugated equine) estrogen did not protect against CVD in men or women.^21^ A randomized controlled trial assessing the cardiovascular safety of testosterone in women has been completed^22^ but not published. The first trial designed to assess the effect of testosterone on major adverse cardiac events in men (NCT03518034) has just started. Regulators in North America have recently warned of cardiovascular risk of testosterone administration.^23 24^ Correspondingly, genetic validation of testosterone as a target of intervention in IHD has recently been demonstrated.^25^ Regulation of the reproductive axis may be more relevant to IHD in men than women because estrogen levels are similar in men and post-menopausal women, but androgen levels are substantially higher in men than women.^26^ As such, GnRH1 might be more likely to affect IHD via testosterone, and be particularly relevant for men. Higher rates of IHD in men than women are not explained by current risk factors.^1^

From a practical perspective this study suggests that drivers and consequences of GnRH1 might be targets of intervention for IHD. Drivers of GnRH1 include energy balance indicated by insulin.^27^ Insulin increases IHD risk.^28^ Insulin raises testosterone^29 30^ providing a pathway from GnRH1 to IHD. Other potential drivers of GnRH1 include neurokinins,^31 32^ nitric oxide,^33^ aspartate/glutamate,^34 35^ gamma-aminobutyric acid^36^ and dynorphin.^37^ Notably, a common feature of several successful treatments for IHD is suppression of the male reproductive axis, including spironolactone, a known anti-androgen, statins,^38^ digoxin,^39^ nitric oxide^40^ and some hypertensives.^41^

## Conclusion

This study confirms in humans the central tenet of evolutionary biology that reproduction trades-off against longevity, by showing that GnRH1, specifically, is positively associated with IHD. This study implies that GnRH1, its drivers and its consequences, are potential targets of intervention for the leading cause of global morbidity and mortality.

## Declarations

### Ethical approval and consent to participate

This article does not contain any studies with human participants or animals performed by any of the authors. As such informed consent is not applicable for this study.

Consent for publication: Not applicable

### Availability of data and materials

This study is an analysis of publicly available data. The program used are available on request

### Author Contributions

CMS had the idea. JCMN and CMS conducted the analysis independently and cross-checked. CMS drafted the manuscript. JCMN reviewed the manuscript for intellectual content and approved the final version.

## Acknowledgements

None.

## Abbreviations

CVD: cardiovascular disease
GWAS: genome-wide association study
GnRH: Gonadotropin-releasing hormone
IHD: ischemic heart disease
SNP: single nucleotide polymorphism

